# Altered polyadenylation site usage in SERPINA1 3’UTR in response to cellular stress affects A1AT protein expression

**DOI:** 10.1101/2024.09.13.612749

**Authors:** FNU Jiamutai, Abigail Hatfield, Austin Herbert, Debarati Majumdar, Vijay Shankar, Lela Lackey

## Abstract

Alternative polyadenylation results in different 3’ isoforms of messenger RNA (mRNA) transcripts. Alternative polyadenylation in the 3’ untranslated region (3’UTR) can alter RNA localization, stability and translational efficiency. The *SERPINA1* mRNA has two distinct 3’ UTR isoforms, both of which express the protease inhibitor α-1-antitrypsin (A1AT). A1AT is an acute phase protein that is expressed and secreted from liver hepatocytes and upregulated during inflammation. Low levels of A1AT in the lung contributes to chronic obstructive pulmonary disease, while misfolding of A1AT in the liver contributes to liver cirrhosis. We analyzed the dynamics of alternative polyadenylation during cellular stress by treating the liver cell line HepG2 with the cytokine interleukin 6 (IL-6), ethanol or peroxide. *SERPINA1* is transcriptionally upregulated after IL-6 treatment and has altered polyadenylation, resulting in an increase in long 3’UTR isoforms. We find that the long 3’UTR represses endogenous A1AT protein expression even with high levels of *SERPINA1* mRNA. *SERPINA1* expression and 3’ end processing were not affected by ethanol or peroxide. IL-6-induced changes in transcriptome-wide transcriptional regulation suggest changes to the endoplasmic reticulum and in secretory protein processing. Our data suggest that inflammation influences polyA site choice for *SERPINA1* transcripts, resulting in reduced A1AT protein expression.

## Introduction

To become mature transcripts the majority of human RNAs undergo 3’ end cleavage and polyadenylation (1). The polyA tail is important for RNA stability and translational efficiency (2). Selection of the polyadenylation cleavage site determines the 3’ end of the transcript. In humans, many genes have multiple possible polyA sites and regulate their 3’ end RNA processing to produce alternative polyadenylation (APA) isoforms (3). Alternative polyadenylation can result in messenger RNA (mRNA) transcripts with dramatically different 3’ UTR lengths and regulatory processes. Alternative polyadenylation can also occur in intronic or upstream coding exons, which can result in transcripts with premature stop codons or altered protein products. The mechanisms that control selection of alternative sites are not well understood. The expression of core polyadenylation proteins, accessory RNA binding proteins and RNA transcript structural accessibility can all influence site selection (4–6). In some cases, processed RNA transcripts with longer 3’UTRs may be able to convert to shorter 3’UTRs, altering their regulatory properties, including stability, localization and translational potential (7, 8).

Most documented alternative polyadenylation is based on tissue specificity, but it is also a key part of cellular differentiation (4, 9). Control of alternative polyadenylation is cell-type specific, and pluripotent cells proliferating and differentiating into specific subtypes with less potency can undergo global 3’UTR lengthening or shortening (9). For example, differentiation of cells into placenta trophoblasts is associated with global 3’ UTR shortening. Interestingly, trophoblast alternative polyadenylation genes are enriched in endoplasmic reticulum (ER) regulatory genes, consistent with a differentiated secretory role in the placenta (10). In addition to differentiation, alternative polyadenylation is also associated with a dynamic response to cellular stressors (11, 12). Quiescent T cells converting to a proliferative state undergo alternative polyadenylation of genes associated with organelles and membrane trafficking (13). Alternative polyadenylation, and specifically lengthening of 3’UTRs, is associated with cigarette smoking in white blood cells (14). Alternative polyadenylation changes the 3’UTR sequencing of transcription and can impact protein expression. For example, polyA site choice can be RNA localization, such as in murine myoblasts where transcripts with longer 3’UTRs are more likely to be associated with the endoplasmic reticulum (15).

Alternative polyadenylation in the *SERPINA1* precursor RNA was recently characterized and associated with tissue-specificity, with longer 3’UTR isoforms normally present in liver tissue (16). The longer 3’UTR isoform of *SERPINA1* is also increased in lung tissues from patients with chronic obstructive pulmonary disease (16). *SERPINA1* mRNA produces α-1-antitrypsin (A1AT) protein, which suppresses the immune response by inactivating neutrophil elastase (17). A1AT is primarily translated in hepatocyte cells and secreted into the bloodstream where it can protect lung tissues from an overactive immune response (18). *SERPINA1* is also expressed at low levels in lung tissues and white blood cells with different transcription start sites and 5’UTR splicing isoforms, resulting in complex post-transcriptional regulation (14, 19–23). Despite the importance of A1AT expression in liver tissue, little is known about *SERPINA1* post-transcriptional regulation in this tissue.

*SERPINA1* is an acute phase gene, meaning that it is upregulated under inflammatory conditions due to a variety of cellular stressors (18). A1AT protein is translated from *SERPINA1* mRNA in the endoplasmic reticulum where it is post-translationally modified and secreted from liver cells as part of its normal biological function (18, 24, 25). Misfolding and ER stress of A1AT variants contribute to A1AT-associated disorders and can lead to chronic obstructive pulmonary disease and liver cirrhosis (26–31). A1AT-related diseases are influenced by genetic variation and environmental conditions, like smoking and alcohol use (32–34). We asked whether alternative polyadenylation was a part of dynamic regulation of *SERPINA1* during cellular stress in liver cells and how this affects A1AT protein expression. We tested the impact of three disease-relevant cellular stressors, the inflammatory cytokine interleukin 6 (IL-6), ethanol and peroxide (35–37). We used the HepG2 liver cell line as a model system to study transcriptome-wide RNA expression and alternative polyadenylation during cellular stress, with a focus on *SERPINA1* mRNA. We found that IL-6 exposure altered the regulation of secreted genes and endoplasmic reticulum-associated genes. IL-6 exposure also affected *SERPINA1* specifically, causing an increase in the long 3’UTR isoforms, which produce less A1AT protein. Neither ethanol or peroxide exposure affect *SERPINA1* expression or 3’ end processing, suggesting that not all cellular stressors act through the same regulatory mechanisms.

## Results

### Inflammation induced by IL-6 alters expression and 3’ end RNA processing of *SERPINA1*

Individuals with chronic obstructive pulmonary disease have an increase in long 3’UTR isoforms of *SERPINA1* in lung tissue, potentially due to chronic inflammatory conditions (16). However, *SERPINA1* mRNA and the A1AT protein product are primarily expressed in the liver (18). We analyzed the 3’UTR of *SERPINA1* from published primary hepatocytes treated with IL-6. The available data is not 3’end specific sequencing, thus it does not accurately quantify polyadenylation sites (38). Analysis of the 3’UTR of *SERPINA1* shows a non-significant trend toward longer 3’UTR alternative polyadenylation isoforms (Supplemental Figure 1b and c). The HepG2 liver cell line expresses *SERPINA1* mRNA and has elevated levels of *SERPINA1* mRNA in response to IL-6 at a lower level than seen in data from published primary hepatocytes treated with IL-6 (38) (**Figure 1b**, Supplemental Figure 1a). The expression and up-regulation of *SERPINA1* in HepG2 cells suggest that this cell line is a consistent model for understanding *SERPINA1* expression and processing in human liver cells. Thus, to test the impact of inflammation on 3’ end processing we treated HepG2 liver cells with the cytokine IL-6. We verified IL-6 induced inflammatory conditions using qRT-PCR for *FGB* and *IL1R1,* which are upregulated in cells exposed to IL-6 (38). As expected, we saw a significant increase in expression for both *FGB* and *IL1R1* (**Figure 1a**). To look specifically at polyadenylation site choice, we performed 3’ end specific Quant-Seq sequencing. QuantSeq has been validated with total RNA sequencing for both differential gene expression analysis and 3’ end identification (39). In addition, we found that *FGB* and *IL1R1* were significantly upregulated in the RNA-seq results after IL-6 treatment, reflecting our qRT-PCR results (Supplementary Figure 1d and e).

**Figure 1.**
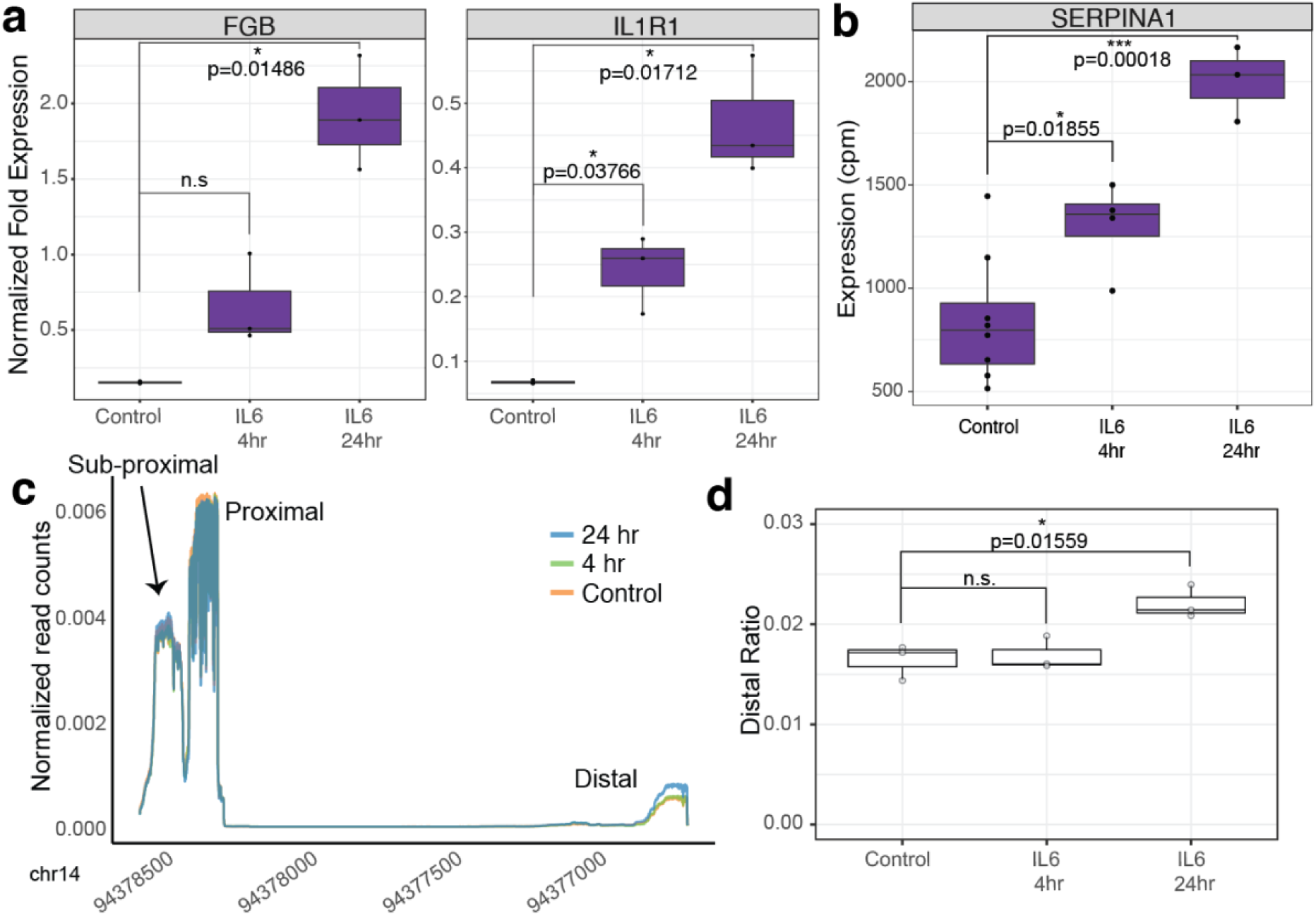
IL-6 induced inflammation changes *SERPINA1* RNA expression and processing in HepG2 liver cells. (a) IL-6 treatment at both 4 and 24 hours results in increased levels of *FGB* and *IL1R1* mRNAs as measured by qRT-PCR (b) and increased expression of *SERPINA1* mRNA as measured by RNA-seq (cpm). (c) 3’ end specific sequencing identifies the majority of *SERPINA1* polyadenylation at the proximal peak. Additional APA occurs at a distal site. IL-6 treatment at 24 hours results in an increase in isoforms using the distal site. Reads are normalized to total reads within the final 3’ exon. (d) Distal *SERPINA1* reads are significantly increased with 24 hours of IL-6 treatment.

Next, we analyzed expression and processing of *SERPINA1* mRNA. We saw significant upregulation of *SERPINA1* at 4 and 24 hours post IL-6 treatment in HepG2 cells (**Figure 1b**). *SERPINA1* is annotated with one polyadenylation site in NCBI RefSeq, corresponding to the long 3’UTR isoform (NM_000295.5) (40). QuantSeq 3’ end specific sequencing shows evidence for both the proximal and distal polyadenylation sites in our HepG2 cell lines, resulting in short and long 3’UTR isoforms respectively (**Figure 1c**). Both the proximal and distal polyA site isoforms are annotated in Ensembl (ENST00000636712.1 and ENST00000393087.9) (41) and have been found in lung and liver tissues (16). We identified a third peak at the very end or within the coding portion of the terminal exon that also corresponds to an annotated transcript in Ensembl (ENST00000449399.7), suggesting that *SERPINA1* may have three polyadenylation sites (**Figure 1c**). We did not find evidence for additional coding or intronic alternative polyadenylation in the HepG2 transcriptome. Treatment of HepG2 cells with IL-6 causes a significant increase in the amount of long *SERPINA1* 3’UTR isoforms 24 hours after IL-6 treatment (**Figure 1c and d**). Processing of *SERPINA1* 3’UTRs was not significantly affected 4 hours post-IL6 exposure.

### IL-6 induced inflammation influences transcriptome-wide expression and 3’ end processing

We identified 827 genes that are altered post-IL-6 exposure (FDR < 0.05 and log_2_FC +/-1) (**Figure 2a and b**). In our transcriptome-wide analysis, *SERPINA1* mRNA is significantly upregulated after 24 hours of IL-6 treatment (**Figure 2b, red**). Approximately half of genes affected by IL-6 treatment at 4 hours remain affected at 24 hours (Supplementary Figure 2a). Overall, more genes are downregulated after IL-6 treatment, especially 24 hours post-IL-6 exposure, however, the median of expression changes is similar in magnitude at both 4- and 24- hour treatments (Supplementary Figure 2b).

**Figure 2.**
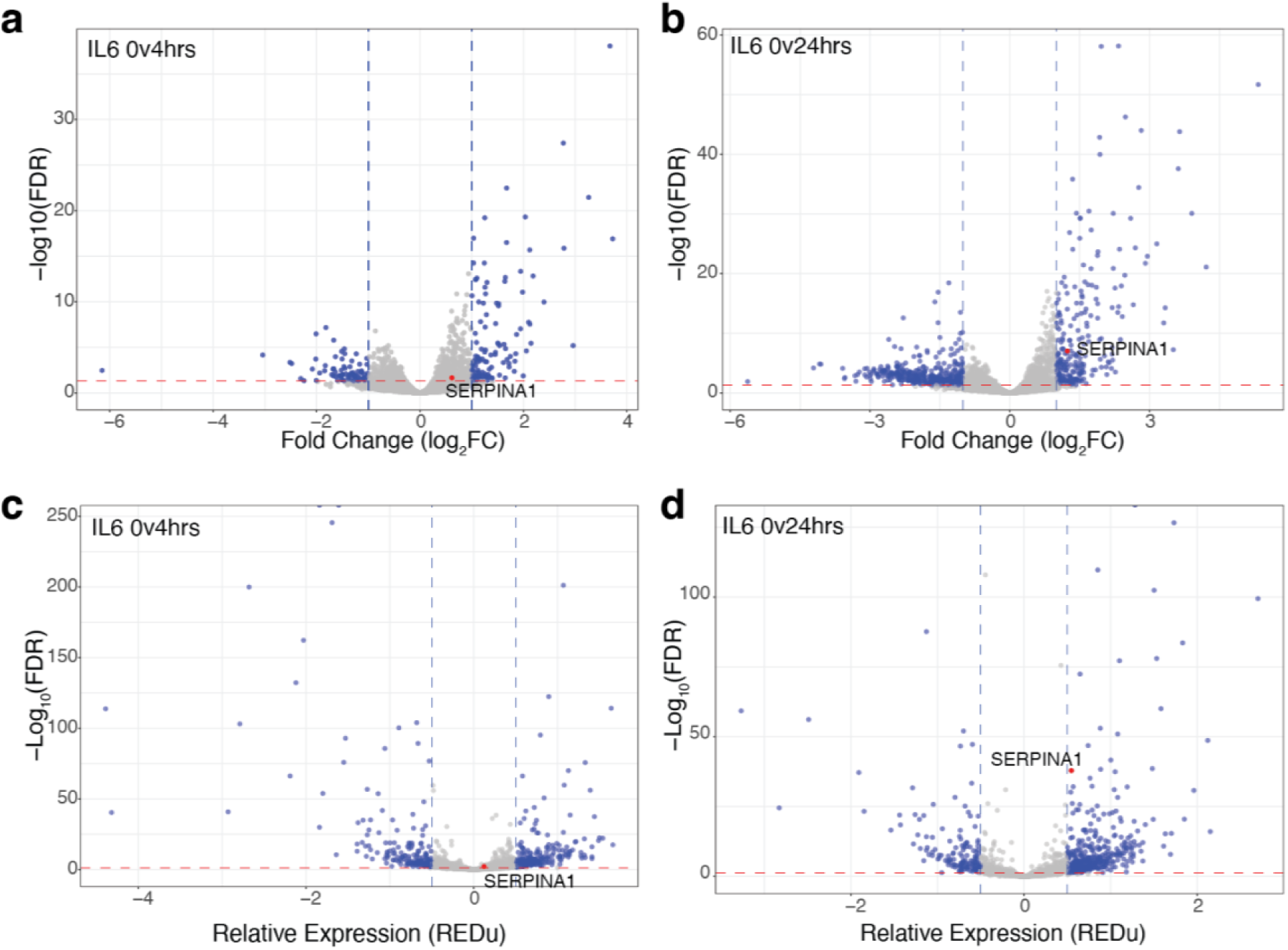
IL-6 induces transcriptome-wide reprogramming, including changes to 3’ RNA processing. Differential gene expression analysis after (a) 4 hours of IL-6 treatment and (b) 24 hours of IL-6 treatment showing transcriptome-wide changes. Alternative polyadenylation analysis with MAAPER after (c) 4 hours of IL-6 treatment and (d) 24 hours of IL-6 treatment showing transcriptome-wide changes. *SERPINA1* expression and significance are highlighted in red.

We used the program MAAPER to quantify alternative polyadenylation transcriptome-wide (42). We found 806 genes with significant changes in polyA site choice at 4 and/or 24 hours post-IL-6 treatment (FDR < 0.05 and REDu polyadenylation score +/- 0.5) (**Figure 2c and d**). MAAPER identified *SERPINA1* as differentially polyadenylated after 24 hours of IL-6 exposure (**Figure 2d**). Approximately half of genes that undergo alternative polyadenylation at 4 hours post-IL-6 treatment remain affected after 24 hours (Supplementary Figure 2c). RNAs become shorter and longer with similar magnitude (Supplementary Figure 2d), however, more genes have longer 3’UTRs after IL-6 exposure. Both polyAdb- and polyAsite-derived polyA site annotations produce similar results for top genes with differential alternative polyadenylation (43, 44) (Supplemental Figure 2e). There is no correlation between RNA levels and alternative polyadenylation in our analysis (Supplemental Figure 2f). Only a small fraction of genes that are differentially expressed are also alternatively polyadenylated (Supplementary Figure 2g).

We found a variety of environmental responses, including the inflammatory and cytokine responses, enriched 4 hours post-IL-6 treatment using gene ontology analysis for biological processes **(Figure 3a)**. Gene list enrichment identified secreted genes and genes associated with protein localization as significantly more likely to be differentially expressed than expected (Human protein atlas - predicted secreted proteins and GO:0008104) (45, 46) **(Figure 3b)**. Metabolic-associated genes were not affected by IL-6 exposure (Human protein atlas - metabolic proteins) **(Figure 3b)**. The top gene ontology pathways shifted at 24 hours post-IL-6 treatment to include cell division and cell cycle pathways **(Figure 3c)**, but secreted genes and protein localization genes remained significantly more likely to be differentially expressed at 24 hours post-IL-6 exposure **(Figure 3d)**. RNA processing genes were significantly depleted from differentially expressed genes at both 4- and 24-hours post-IL-6 treatment (GO:0006396) (46) **(Figure 3b and d)**, suggesting that RNA processing genes are less likely to be differentially expressed than other genes.

**Figure 3.**
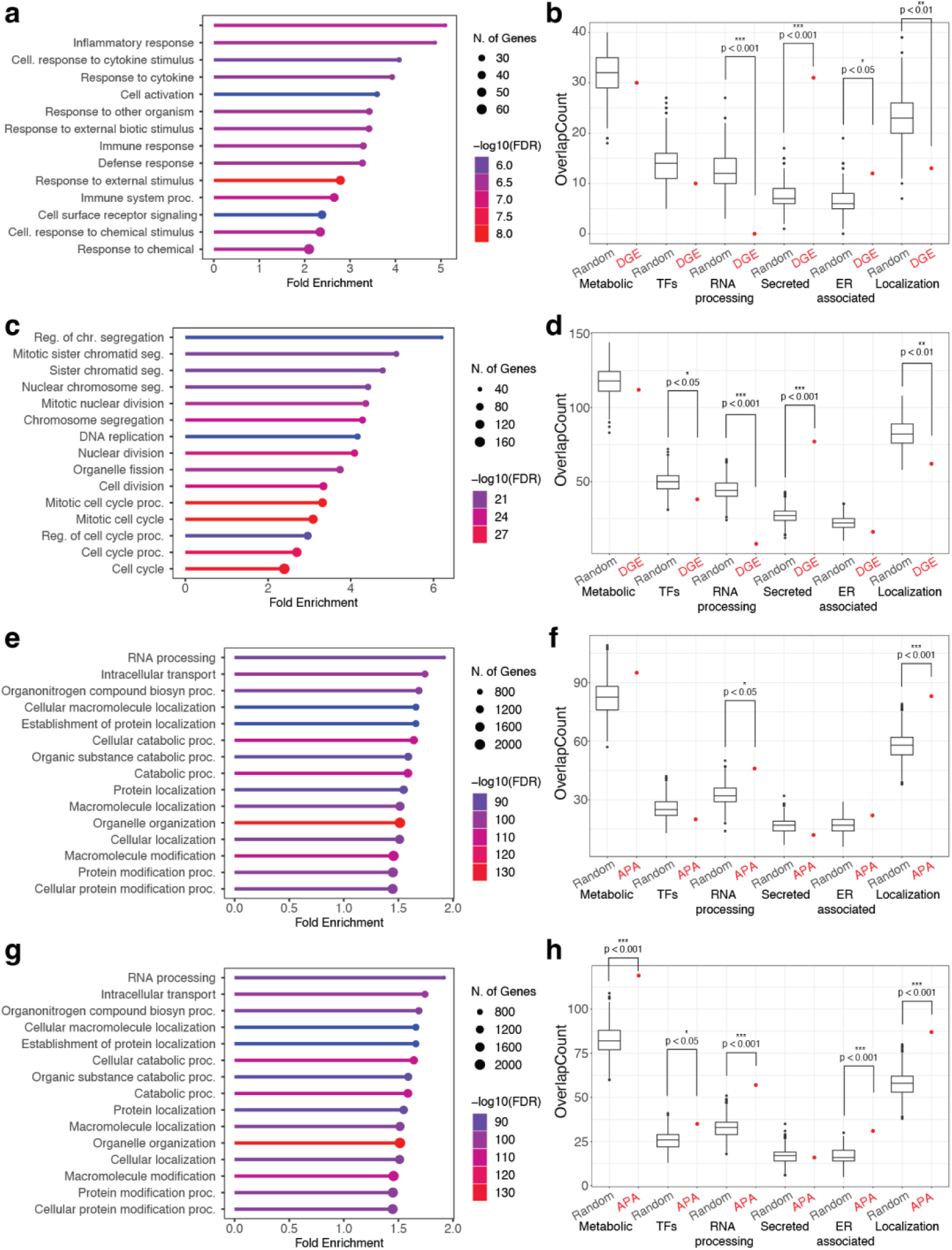
Endoplasmic reticulum genes are differentially expressed and alternatively polyadenylated during the inflammatory response. (a) Response to inflammation and the environment were top biological processes 4 hours post-IL-6 exposure while (b) secreted genes were overrepresented in gene sets. (c) 24 hours post-IL-6 exposure, cell cycle-associated biological processes were enriched and (d) secreted genes continued to be overrepresented. APA analysis (e) 4 hours post-IL-6 exposure gene ontology and (f) gene-set overrepresentation identified RNA processing and protein localization-associated genes. Both RNA processing and protein localization genes were also identified by (g) ontology and (h) enrichment analyses 24 hours post-IL6 exposure.

When we analyzed alternative polyadenylation data using gene ontology, we found enrichment for RNA processing at both 4- and 24-hours post-IL-6 treatment **(Figure 3e and g)**. This enrichment was consistent in gene list analysis where RNA processing was more likely to be differentially expressed compared to random **(Figure 3f and h)**. These results suggest that RNA processing genes are affected by the inflammatory response at the post-transcriptional level but not at the transcriptional level. Genes associated with protein localization were more likely to be differentially expressed at both 4- and 24-hours post-IL6-exposure **(Figure 3f and h)**. After 24 hours of IL-6 exposure genes associated with the endoplasmic reticulum were significantly more likely to be alternatively polyadenylated (Human protein atlas – endoplasmic reticulum) (45) **(Figure 3h)**. Both protein localization and endoplasmic reticulum-associated genes are likely affected by IL-6 through both transcriptional and post-transcriptional regulation, including 3’ processing. Inflammation-induced alternative polyadenylation of endoplasmic reticulum-associated genes is only evident after 24 hours of exposure, suggesting a delayed response **(Figure 3h)**. Previous studies have associated alternative polyadenylation with regulation of secreted proteins, such as within B cells and placental cells (10, 15). Our results indicate that liver cells may alter their secretory network during inflammation by controlling RNA levels and regulating RNA 3’ end processing.

### 3’ end RNA processing regulation in *SERPINA1* precursor mRNA affects A1AT protein expression

*SERPINA1* produces A1AT, which is secreted from liver hepatocytes. Previous research showed that the long 3’UTR isoform of *SERPINA1* was associated with repression of protein synthesis using nanoluciferase reporter assays (16). We developed a *SERPINA1* CRISPR mutant cell line to determine whether 3’ end processing influences A1AT protein expression from the endogenous *SERPINA1* locus expressed in HepG2 liver cells (APA-mut line) (**Figure 4a**). Treatment of wild-type and APA-mut HepG2 cells with IL-6 results in similar overall gene expression changes (Supplemental Figure 3a-c). However, *SERPINA1* mRNA has higher expression in the APA-mut cell line than in wildtype HepG2s both before and after IL-6 treatment (**Figure 4b**, Supplemental Figure 3a-c, red points). APA-mut lines dramatically shift from predominately proximal polyadenylation of *SERPINA1* to predominantly distal polyA sites (**Figure 4c**). Although the main proximal *SERPINA1* site is mutated and unable to be used for processing, we see rarely used post-proximal and sub-distal alternative polyadenylation sites that generate novel shorter 3’UTRs (**Figure 4c**). Although *SERPINA1* mRNA is expressed at overall higher levels in APA-mut cells, IL-6 treatment induces a similar up-regulation of *SERPINA1* mRNA in both lines (**Figure 4b**, Supplemental Figure 3d and e, red points). However, IL-6 treatment of APA-mut cell lines does not increase the distal polyA site usage as does in wild-type HepG2 cells (**Figure 4d**). Consistent with this, MAAPER transcriptome-wide analysis does not identify *SERPINA1* as alternatively polyadenylated after IL-6 treatment. The loss of IL-6 mediated 3’ processing when the proximal site is mutated implies that the increase in distal polyadenylation is controlled by regulation of the proximal polyA site.

**Figure 4.**
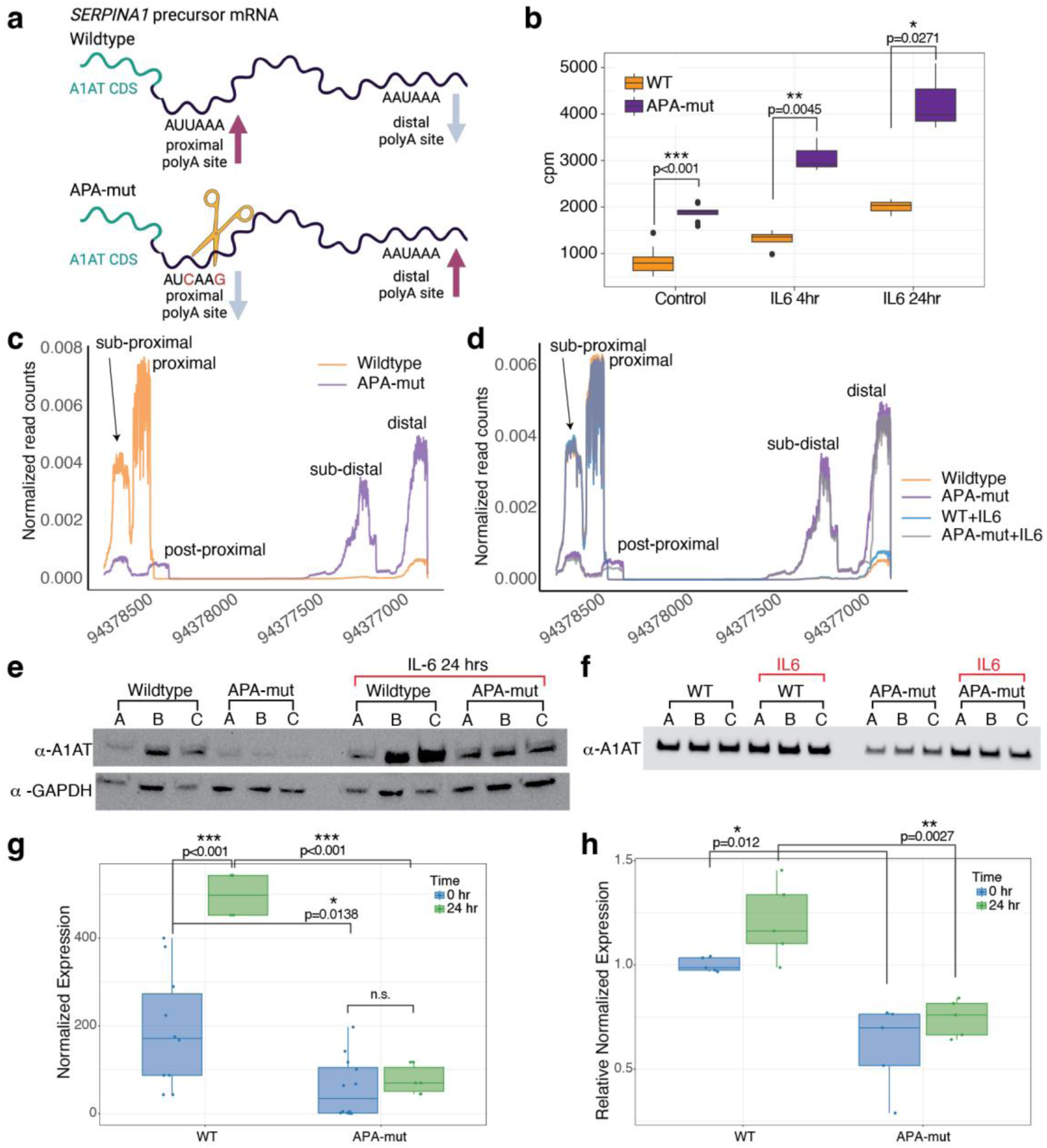
The long 3’UTR of *SERPINA1* represses A1AT protein expression. (a) APA-mut HepG2 cells were created by CRISPR mutation of the proximal polyA site from AAUAAA to AUCAG (Synthego). (b) APA-mut cells express significantly more *SERPINA1* mRNA than wild-type and increase expression upon IL-6 treatment. (c) Mutation of the proximal polyA site in the APA-mut cell line (purple) increases the use of distal polyA sites. (d) Treatment of wildtype HepG2 cells results in an increase in distal polyA usage after 24 hours, but does not affect the ratio of distal polyA site use in APA-mut cells. (e,g) A1AT protein expression is decreased in the APA-mut cell line. A1AT protein expression is upregulated after IL-6 treatment, but upregulation remains impaired in the APA- mut cell line. (f,h) Secreted A1AT protein is decreased in the APA-mut cell line. IL-6 treatment does not significantly increase secreted levels of A1AT.

While differential gene expression is consistent between wild-type and APA-mut lines at 4- and 24-hours post-IL-6 exposure, alternative polyadenylation is not highly correlated 4 hours post-IL-6 exposure (Supplemental Figure 3f). Instead we only see high correlation 24 hours post-IL-6 exposure, suggesting that alternative polyadenylation may be a longer-term regulation for IL-6 mediated inflammation in HepG2 cells (Supplemental Figure 3g).

Reporter assays have shown that long 3’UTR isoforms of *SERPINA1* repress nanoluciferase protein expression (16). Therefore, we tested A1AT protein production in the wildtype and APA-mut cell lines. Despite the higher levels of *SERPINA1* mRNA in APA-mut cells, A1AT protein levels were much lower in mutant cells, which primarily produce the long 3’UTR isoform of *SERPINA1* (**Figure 4e**, Supplemental Figure 4). A1AT protein expression is significantly upregulated after 24 hours of IL-6 exposure (**Figure 4e**, Supplemental Figure 5). A1AT protein in the mutant line is not significantly upregulated after IL-6 treatment and remains at a much lower level than in wildtype HepG2 cells (**Figure 4e and g)**. Quantification of protein and RNA levels shows that 24 hours of IL-6 exposure induces *SERPINA1* mRNA upregulation by 1.24-fold, while A1AT protein increases by 2.62-fold. In contrast, in APA-mut cells, while IL-6 exposure increases *SERPINA1* mRNA by a similar amount (1.18-fold), A1AT protein levels only increase by 1.32-fold (**Figure 4g**). Since A1AT is secreted from hepatocytes, we tested the levels of A1AT protein in the media. We found significantly more A1AT secreted from wildtype HepG2 cells than from APA- mut HepG2 cells (**Figure 4f and h**). IL-6 exposure showed a minor but non-significant increase for both wildtype and mutant cells (**Figure 4f and h**), indicating that changes in *SERPINA1* mRNA levels and intracellular A1AT expression may not directly correlate with secreted A1AT.

### SERPINA1 is not affected by the cellular stress induced by ethanol and peroxide exposure

Liver stress can be caused by ethanol exposure (47, 48). Ethanol induces reactive oxygen species (ROS) (37). To test the impact of acute ethanol exposure on 3’ end processing we treated cells with 170 mM or 300 mM ethanol for 3 or 24 hours. We used qRT-PCR to analyze the effect of ethanol treatment on *BCL2* and *GPX2* mRNAs. When we treated cells with 300 mM ethanol, we saw a significant response to *BCL2* after 4 hours of ethanol treatment and to *GPX2* at 24 hours of ethanol exposure (**Figure 5a**). This aligns with published literature on the *BCL2* and *GPX2* response to ROS (49, 50). We did not find a significant response to ethanol treatment with 170 mM ethanol, although the expression trend is similar between the two conditions. We used cells treated with 170 mM ethanol for 3’ end specific library preparation and sequencing to avoid ethanol-induced toxicity. RNA-seq data for *GPX2* are consistent with qRT-PCR data, with a non-significant trend in increased expression with ethanol treatment (**Figure 5b**). In addition to ethanol treatment, we also treated cells with peroxide directly to analyze the ROS pathway separate from other effects caused by ethanol exposure. *GPX2* is significantly upregulated with peroxide treatment, as expected (49). *SERPINA1* expression is not significantly affected by ethanol or peroxide treatment (**Figure 5c**).

**Figure 5.**
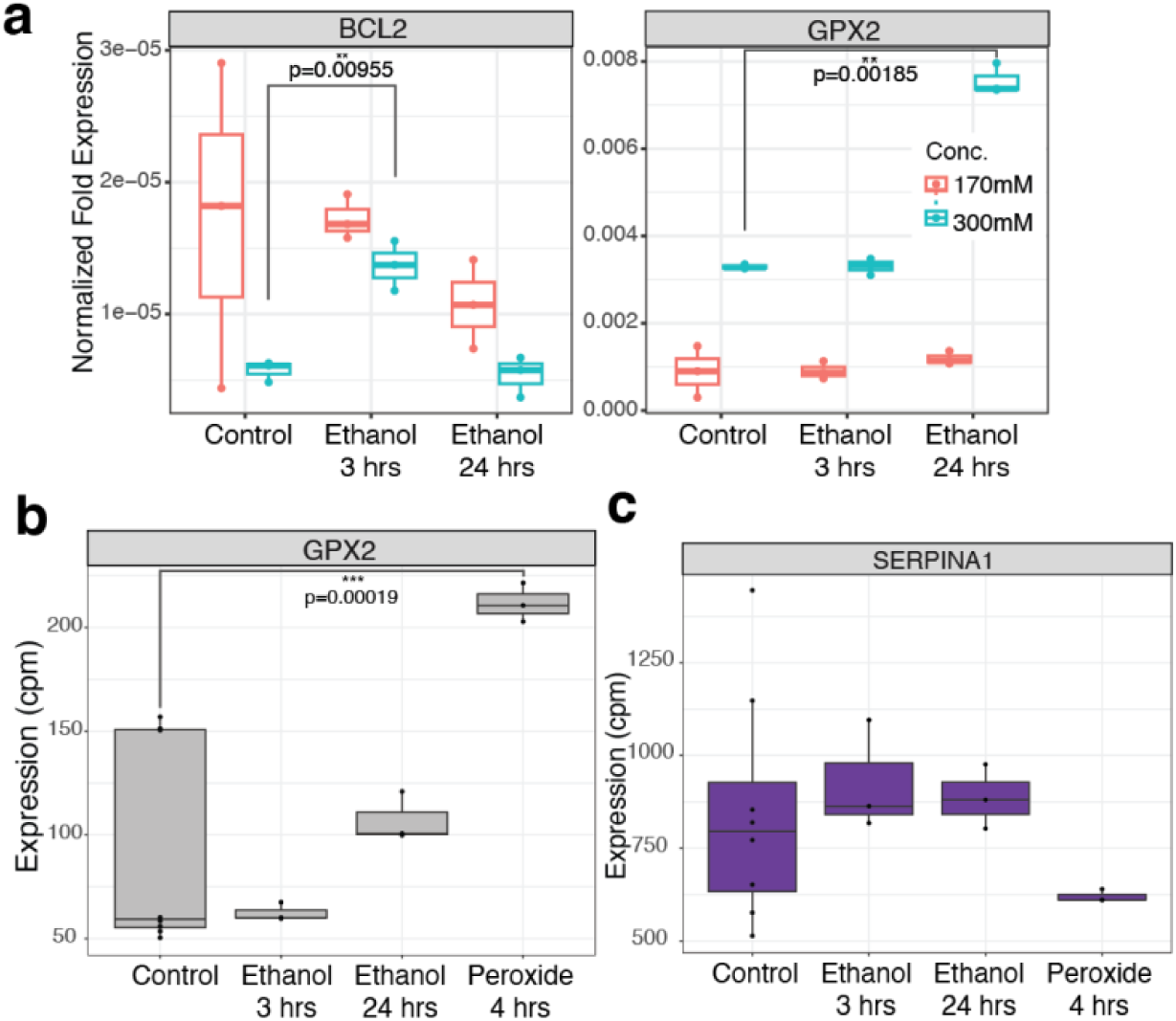
Cellular response to ethanol and peroxide does not alter *SERPINA1* mRNA expression. (a) *BCL2* and *GPX2* have moderate expression changes with ethanol treatment as quantified by qRT-PCR. (b) RNA-seq data follows the same non-significant trend for *GPX2* expression with 170mM ethanol treatment, but shows upregulation of *GPX2* with peroxide exposure. (c) RNA-seq data of *SERPINA1* mRNA expression shows similar levels of mRNA at both 3 and 24 hours post-ethanol exposure and with peroxide treatment.

Ethanol caused mild perturbation to the transcriptome with 96 and 26 differentially expressed genes at 3 and 24 hours respectively (FDR < 0.05 and log_2_FC +/-1) (**Figure 6a and b**). Only a small number of genes are differentially expressed at both 3 and 24 hours (Supplemental Figure 7a), however, these genes are consistently up or down regulated (Supplemental Figure 7b, log_2_FC +/-0.5). In contrast, our peroxide treatment was harsher and had a major impact on the transcriptome, with 572 genes differentially expressed after 4 hours of treatment (FDR < 0.05 and log_2_FC +/-1) (**Figure 6c**). Ethanol and peroxide treatment have minimal overlap in the number of differentially expressed genes (Supplemental Figure 7c).

**Figure 6.**
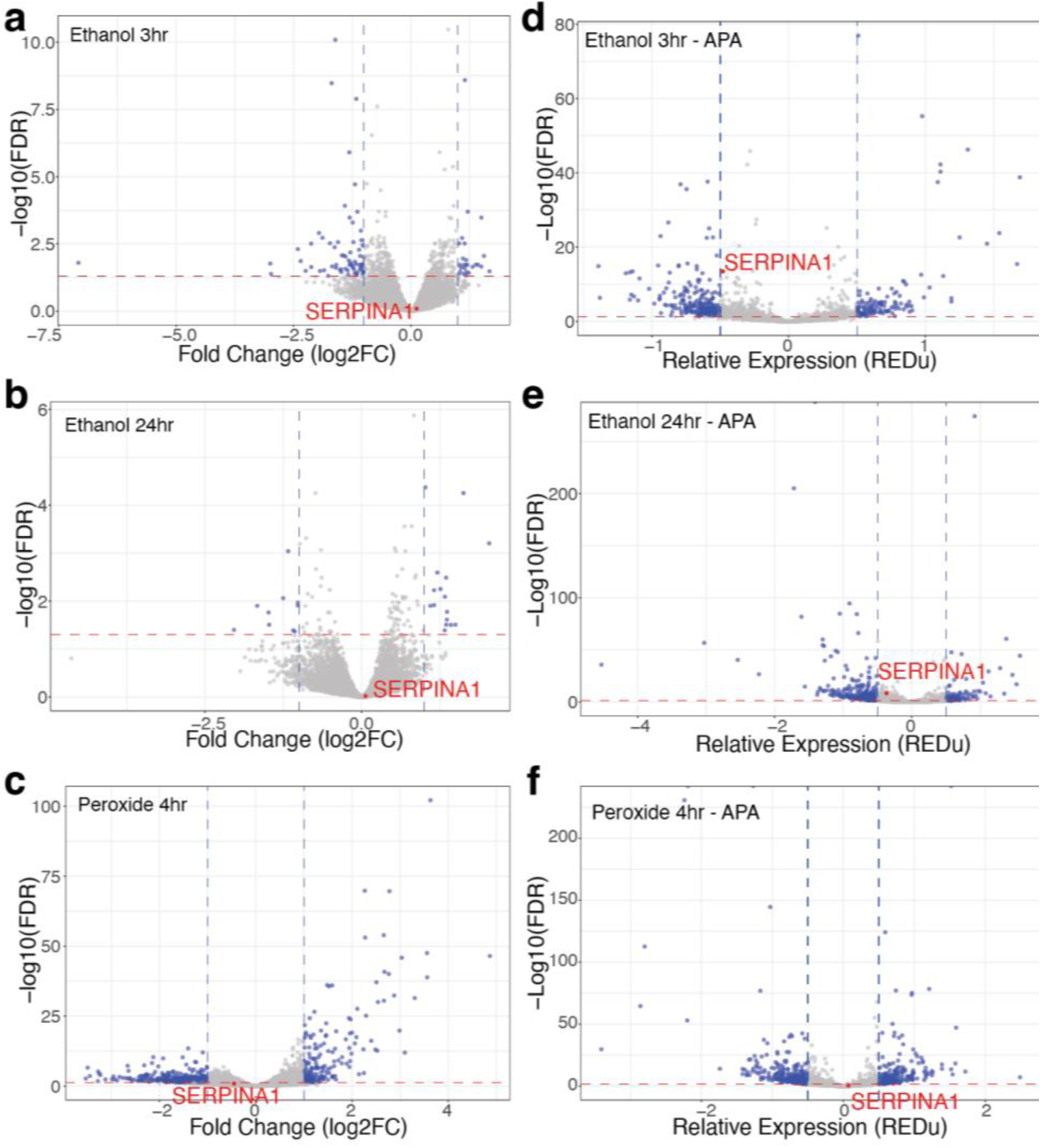
Ethanol and peroxide exposure do not affect *SERPINA1* expression or 3’ end processing. Ethanol treatment at 170mM for (a) 3 hours or (b) 24 hours alters gene expression as measured by RNA levels. *SERPINA1* is unaffected (red). (c) Peroxide exposure over four hours affects gene expression transcriptome-wide, but does not significantly alter *SERPINA1* mRNA levels (red). (d) Transcriptome-wide alternative polyadenylation is altered following (d) 3 hours or (e) 24 hours of 170mM ethanol treatment or (f) 4 hours of peroxide exposure. *SERPINA1* mRNA alternative polyadenylation is not significantly impacted with ethanol or peroxide treatments (red).

*SERPINA1* 3’ end processing is not altered by ethanol or peroxide exposure (**Figure 6d-f**). However, both ethanol and peroxide impact RNA 3’ processing transcriptome-wide, with ethanol affecting alternative polyadenylation for 581 genes (**Figure 6d and e**) and peroxide affecting alternative polyadenylation for 530 genes (**Figure 6f**) (FDR < 0.05 and REDu +/-0.5). Approximately a third of genes with differential polyadenylation 3 hours post-ethanol exposure remain differentially polyadenylated at 24 hours post-ethanol exposure (Supplemental Figure 7d). Some genes that are alternatively polyadenylated by ethanol treatment are also alternatively polyadenylated with peroxide exposure (Supplemental Figure 7e). Ethanol and peroxide have a minor impact on the endoplasmic reticulum and secreted genes although the response is not as consistent or strong as IL-6 exposure (Supplemental Figure 8a-f).

## Discussion

Alternative polyadenylation has been linked to secretory pathways during the differentiation of placental secretory cells and the conversion of B cells to antibody secreted plasma cells (10). Liver hepatocytes have an important secretory role, producing high levels of serum proteins (51). Inflammation induces upregulation of acute phase proteins like A1AT, which must be synthesized and secreted from the liver (51). In our study, differentially expressed and alternatively polyadenylated genes impact endoplasmic reticulum processes. We also find that secreted genes themselves are regulated at the RNA level. A subset of genes have longer 3’UTRs after IL-6 exposure, including *SERPINA1*. Further analysis of these genes to identify co-regulatory mechanisms is important for understanding inflammatory RNA regulation.

Although *SERPINA1* mRNA levels increase after IL-6 exposure, like most acute phase genes (Supplemental Figure 9a), we did not find most acute phase genes to have altered polyA site choice using MAAPER transcriptome-wide. Since we manually modified the MAAPER polyAdb annotations to encompass known 3’ ends for *SERPINA1,* we directly analyzed 3’UTR reads in all acute phase genes expressed in HepG2 cells. This analysis identified additional upregulated acute phase genes, ceruloplasmin (*CP)* and haptoglobin (*HP),* as alternatively polyadenylated (Supplemental Figure 9b). While the polyA choice for most acute phase genes remains the same with IL-6 exposure (11/14 genes) (Supplemental Figure 9c), our analysis does show that alternative polyadenylation regulates a substantial fraction of acute phase genes during inflammation (∼20%). However, the identification of *CP* and *HP* by manual analysis suggests that additional work is needed to develop comprehensive 3’ end annotation sets for alternative polyadenylation analysis.

As human liver is a difficult system to study, we used the HepG2 cell line as a model system. We confirmed that HepG2 cells behave similarly to primary human hepatocytes with IL-6 exposure with upregulation of *SERPINA1* (*38*). In addition, we have previously confirmed the existence of *SERPINA1* long and short 3’UTR isoforms in primary liver and lung tissues (16). Future studies may be able to more directly access human tissues and study alternative polyadenylation in an more complex or primary context. For example, treatment of HepG2 cells with chemical stressors cannot replicate the environment of the human liver where immune cells co-exist with hepatocytes and supporting liver cell types (52). While we did not find an impact of ethanol or peroxide treatment on *SERPINA1* expression or 3’ end processing, this may be due to our short-term, simplified treatment model. A better understanding of the impact of ethanol on 3’ end processing and *SERPINA1* regulation in hepatocytes may require 3D organoid systems, a chronic ethanol exposure time-course or co-exposure of ethanol with the pro-apoptotic cytokine TGF-ß (53).

In *SERPINA1,* mutation of the proximal polyA site prevents IL-6 mediated 3’UTR regulation. This suggests that the proximal polyA site is the primary regulatory target and the increase in long 3’UTR isoforms results from repression of the proximal polyA site. The *SERPINA1* long 3’UTR isoform is less capable of producing A1AT protein. Our data suggest that while *SERPINA1* mRNAs are upregulated during the early inflammatory response, A1AT protein production is muted by non-productive long 3’UTR transcripts. This allows the cells to initiate high levels of RNA transcription but reduced translation of A1AT and potentially alleviate an overburdened secretory system. In addition, long 3’UTR transcripts are capable of conversion to short 3’UTR isoforms, suggesting that *SERPINA1* long 3’UTRs may be converted to productive transcripts at later stages or through other regulatory processes (7, 8).

The mechanism underlying *SERPINA1* long 3’UTR isoform repression remains unknown, but several RNA binding proteins have been linked to A1AT repression, including QKI and NQO1 (16, 54). In addition, the miR-320 family has been implicated in translational regulation of A1AT expression during inflammation (55). Exogenous A1AT protein represses endogenous production of *SERPINA1*, suggesting the existence of a negative feedback loop and highlighting the importance of both transcriptional and post-transcriptional control of *SERPINA1* translation of A1AT (56). Additionally, upregulation of A1AT protein after IL-6 surpasses the upregulation of the mRNA, implying that post-transcriptional regulation may also act on the shorter 3’UTR isoforms of SERPINA1 to control A1AT translation.

A1AT deficiency is linked to a rare variant that causes A1AT misfolding and endoplasmic reticulum stress in the liver (31). Our work identifies how inflammation and other cellular stressors influence gene expression and 3’ end processing of *SERPINA1* and other genes. We find that inflammation changes transcriptome-wide RNA expression and 3’ end processing for genes associated with the endoplasmic reticulum and protein localization. Remodeling of these secretory pathways during inflammation may contribute to disease and provide therapeutic targets to alleviate specific pathology, such as the role of A1AT misfolding and accumulation in hepatocytes in liver cirrhosis. Future studies on mechanism are necessary to understand how inflammation causes alternative polyadenylation of endoplasmic reticulum and protein localization genes and the functional impact of these APA events.

## Methods

### Cell culture

APA-mut cell lines were created through Synthego by CRISPR mediated mutation of AUUAAA to AUCAAG on the minus strand of chr14:94,378,388-94,378,393 (hg38 genome). HepG2 wildtype cells were also provided from Synthego and regularly checked for mycoplasma contamination (PCR Mycoplasma Detection Kit, ABM). Cell lines were seeded in 10 cm^2^ plates and grown at 5% CO^2^ and 37 C. Cells were grown in Eagle’s Minimum Essential Media (EMEM) supplemented with 5% FBS (Thermo Scientific) and 0.5% Penicillin-Streptomycin (MilliporeSigma). For exposure treatments, cell lines were seeded into at 3.0 x 10^5 cells in 6- well plates. Cells were incubated for 24 hours then serum-starved using 1 mL serum-free EMEM for another 24 hours. After media starvation, the cells were treated in 1 mL serum-free media with 170mM ethanol diluted in PBS (EtOH, 3, and 24 hours. Fisher Scientific), 150 uM hydrogen peroxide diluted in PBS (H2O2, 4 hours. Fisher Scientific) or 20ng/mL interleukin-6 diluted in 0.1% of BSA at (IL-6, 4, and 24 hours. Fisher Scientific).

### RNA extraction

Total RNA was extracted using Trizol (Thermo Fisher) according to manufacturer’s recommendations for adherent cell lines using Phaselock columns (Quantabio) prior to RNA purification with RNA clean up columns (NEB). Genomic DNA was removed using TURBO DNase (Fisher Scientific) according to the given protocol. RNA was quantified via NanoDrop (Thermo Scientific ND-8000) and quality checked by RNA Screentape on a Tapestation 4150 (Agilent).

### Quantitative reverse transcriptase polymerase chain reaction (qRT-PCR)

qRT-PCR was performed on a QuantStudio 3 (Thermo Scientific) system using iTaq Universal SYBR Green One-Step Kit (Biorad). 150 ng of total RNA was loaded, in biological triplicate, into a total reaction of 20 uL. qRT-PCR run parameters were as follow: 10 minutes reverse transcription at 50 C, 1 minute polymerase activation and DNA denaturation at 95 C, followed by 40 cycles involving denaturation at 95 C for 15 seconds, annealing/extension/plate read at 60 C for 1 minute, and a melt curve analysis (95 C, 15 sec; 60 C, 1 min; 95 C, 1 sec with 0.15C /s ramp rate). The sequences for the primers used for genes *BCL2, GPX2, GAPDH, ACTB, FGB,* and *IL1R1* are provided (Supplemental Table 1). At least three biological replicates were quantified and a t-test was used to determine statistical significance.

### RNA library preparation and sequencing

RNA samples were prepared using the QuantSeq 3’ mRNA REV Library Prep Kit V2 (Lexogen), which captures polyadenylated RNA. Quality assurance for loading samples for sequencing was performed using a Qubit dsDNA High Sensitivity Assay (Thermo Scientific) on a Qubit 4 fluorometer (Thermo Scientific), and a High Sensitivity D1000 Screentape (Agilent) on a Tapestation 4150 (Agilent). The Lexogen QuantSeq samples were sequenced on a NovaSeq 6000 (Illumina) using a 2×100 cycle cartridge. A custom sequence primer was used for Read 1 according to instructions from Lexogen and Illumina.

### Quantification of APA events

Direct coverage of select regions were conducted through Samtools mpileup counts (57). Each position’s normalized read count was calculated by dividing total read counts of region from the read counts of each individual position. To calculate the length normalized ratio of reads within distal and proximal sections of the 3’UTR we divided the read counts by the length of the region and then divided this length normalized value by the sum of the read ratios of all regions. To quantify transcriptome-wide APA we used the R package MAAPER (42) with read length of 100 and num_pas_thre of 50. We modified the polyAdb annotations (43) provided with MAAPER to include the APA sites originally unannotated in *SERPINA1.* In addition, we used MAAPER with annotations from the polyAsite database (44). To adjust the significance for multiple testing we used Benjamini and Hochberg’s False Discovery Rate correction (58). Unless otherwise stated, differential polyadenylation was determined as genes with FDR < 0.05 and polyA score REDu greater or less than 0.5. Differentially polyadenylated genes were used as input into shinyGO for biological pathway analysis (59). Differentially polyadenylated genes were used to compare against randomly selected HepG2 expressed genes using bootstrapping analysis for enrichment in gene lists from amiGO (46) and the Human Protein Atlas (proteinatlas.org) (45). Permutation tests were used to determine statistical significance and reported as p-value < 0.05, 0.01 or 0.001.

### Differential gene expression analysis

For differential gene expression analysis we used an edgeR-based pipeline for TMM normalization and quantification (available on GitHub https://github.com/vshanka23/snakemake_rnaseq), and expression modeling (“RNA-seq with edgeR”, https://github.com/bmunn99/Clemson-Grad-Rscripts). The edgeR package was used to normalize library sizes and quantify expression levels for individual genes (60). Differentially expressed genes passed the following filters unless otherwise stated: FDR < 0.05, log_2_CPM > 6 and log_2_FC > 1. Differentially expressed genes were used as input into shinyGO for biological pathway analysis (59). Differentially expressed genes were used to compare against randomly selected HepG2 expressed genes using bootstrapping analysis for enrichment in gene lists from amiGO (46) and the Human Protein Atlas (proteinatlas.org) (45). Permutation tests were used to determine statistical significance and reported as p-value < 0.05, 0.01 or 0.001.

### Western blot analysis

HepG2 cells were harvested using TrypLE, spun down at 500 x g, then resuspended in 500uL NP-40 buffer, with the addition of HALT Protease Inhibitor (Thermo Scientific). Column concentrators were used for media concentration which was volume normalized (Pierce Concentrator PES 10K MWCO). Cells were lysed for 20 minutes on ice. Lysis was reduced and denatured using 4X BOLT LDS Sample Buffer (Novex). Samples were vortexed then denatured at 70°C for 10 minutes then run on Bolt 10% Bis-Tris Plus Wedgewell Gels (Invitrogen) in 1X MES SDS Running Buffer (Invitrogen). Semi dry transfer was performed using the iBlot2 Gel Transfer Device (Invitrogen) with Iblot 2 NC Mini Stack (Invitrogen). Membrane was stained with a Ponceau S solution (Sigma) and imaged for total protein load on a Chemidoc MP (Biorad). The Ponceau S was removed (0.1% NaOH in nuclease-free water) and the membrane blocked for one hour with (5% fat-free milk, 1X TBS-T buffer). Blots were incubated with A1AT antibody (ThermoFisher #MA5-14661, 1:1000) for two hours then washed with TBS-T. Membranes were incubated with anti-mouse IgG HRP linked secondary (Cell Signaling Technology #7076, 1:2000) for one hour. This procedure was repeated for GAPDH (Santa Cruz #SC-32233, 1:2000) with the same secondary. The blots were washed then incubated in Clarity Western ECL substrate (BioRad) for five minutes. The proteins of interest were imaged using a Chemidoc MP on the Chemiluminescent setting with appropriate exposure times. Protein band quantification was performed using Image Lab software and signals were normalized to GAPDH or Ponceau S protein expression. At least three biological replicates were quantified and a t-test used to determine statistical significance.

## Author contributions

AbH and J performed the experiments. J, AuH, VS, DM and LL performed the bioinformatics analysis. J and LL wrote the manuscript. All authors approved the final manuscript.

## Data availability statement

All RNA-seq data will be submitted to the Sequence Read Archive (SRA): BioProject XXX. Additional datasets used for comparative analysis are publicly available from the Gene Expression Omnibus (GEO) GSE202045.

## Additional Information (including a Competing Interests Statement)

The authors declare no competing interests.

## Supplemental Figures

**Supplemental Figure 1.**
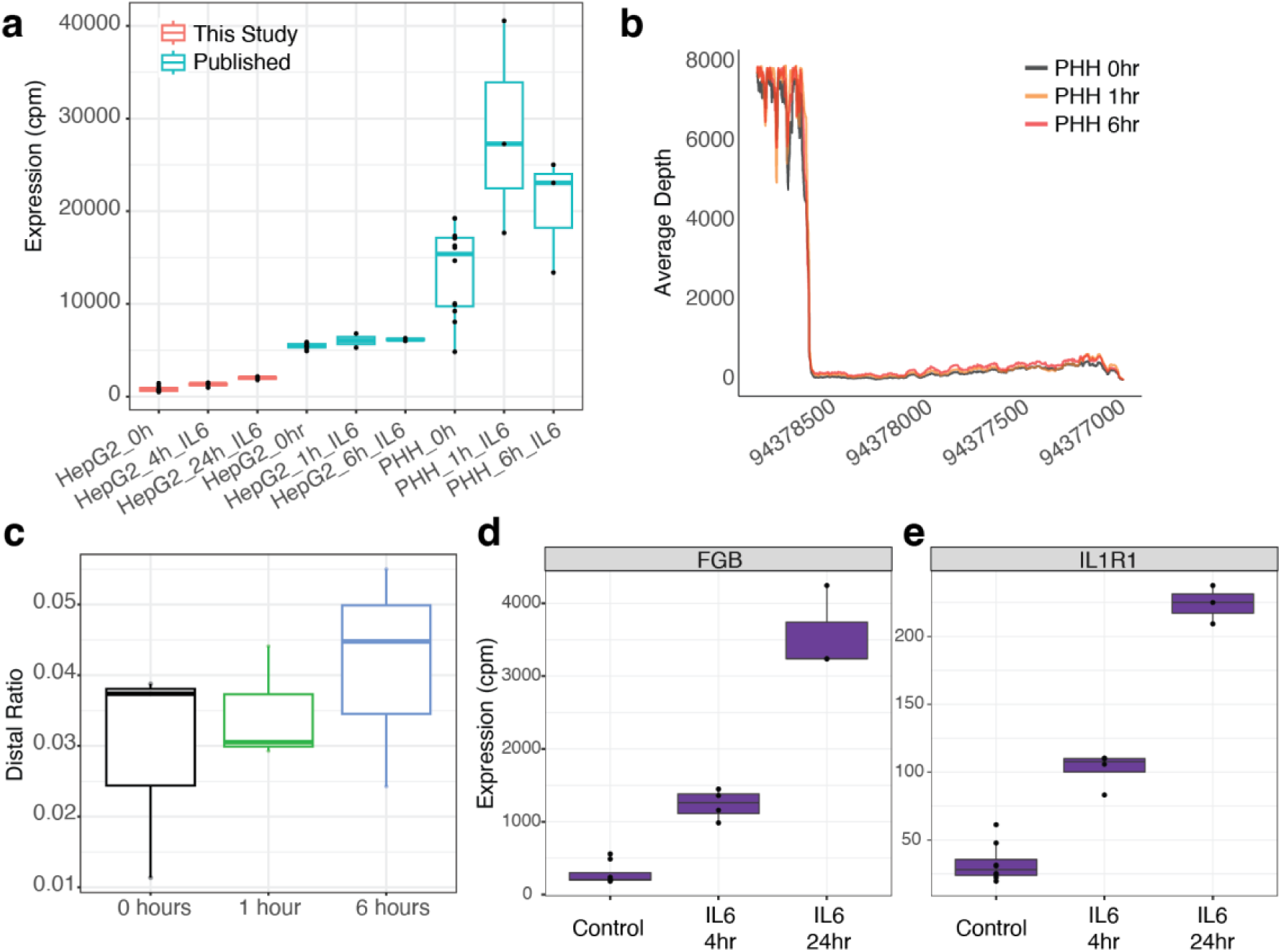
Expression and processing in HepG2 and primary human hepatocytes of *SERPINA1*, *FGB* and *IL1R1.* (a) Increased *SERPINA1* expression in HepG2 cells before and after 4 hrs or 24 hrs of IL-6 exposure (this study), *SERPINA1* expression in HepG2 cells and primary human hepatocytes (PHH) before and after 1 and 6 hrs of IL-6 exposure (published, GSE202045). 3’ end processing in *SERPINA1* in primary human hepatocytes (b) read depth from RNA-seq data and (c) quantification by proximal and distal regions does not identify clear differences before and after IL-6 treatment. Upregulation of (d) *FGB* and (e) *IL1R1* after IL- 6 treatment of HepG2 in Quant-Seq data is consistent with qRT-PCR quantification.

**Supplemental Figure 2.**
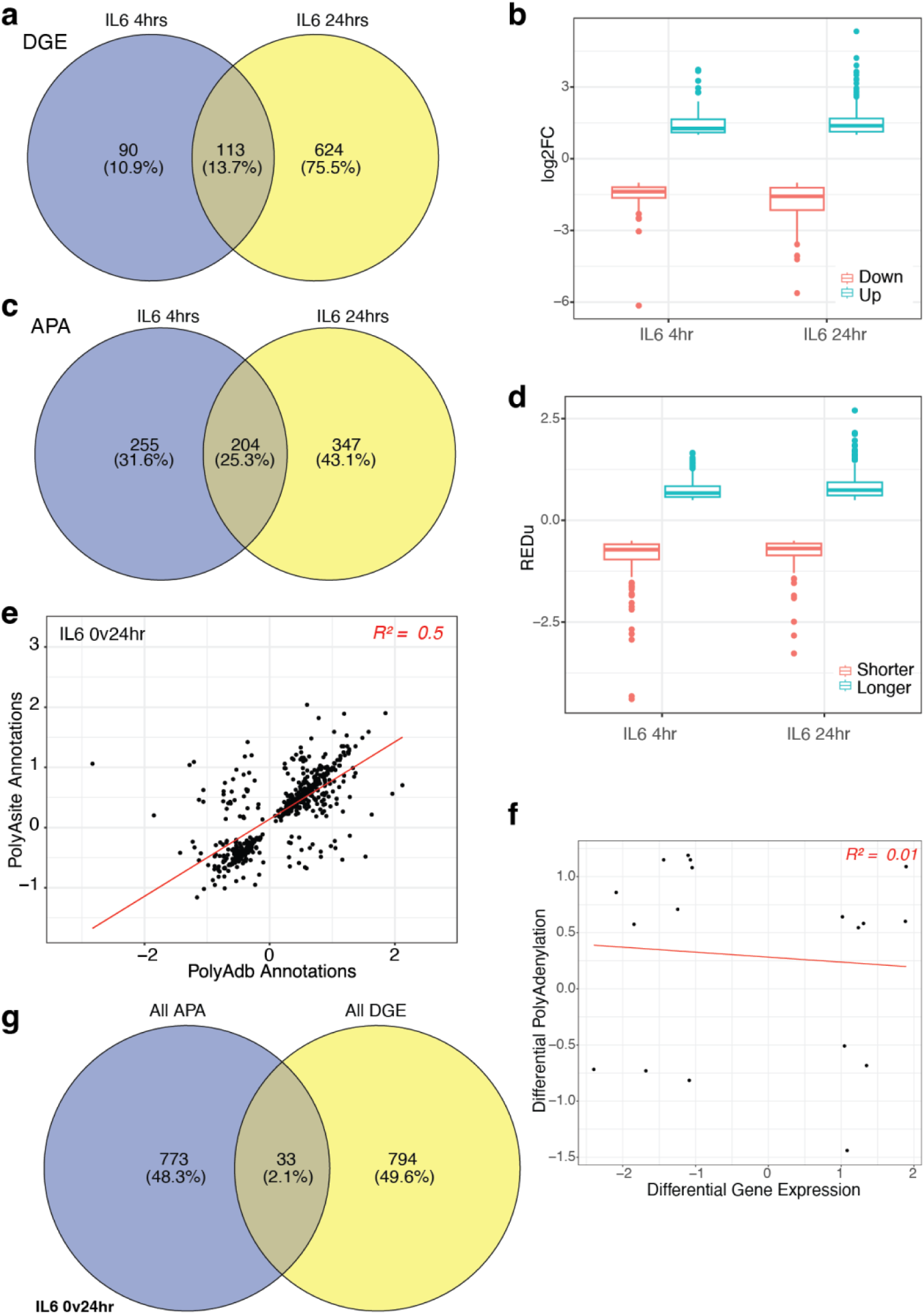
Properties of genes that undergo IL-6 induced differential expression and alternative polyadenylation. (a) Overlap in genes identified as differentially expressed (DGE) at 4hr and 24hr post-IL-6 treatment and (b) the extent of up or down-regulation at each time point. (c) Overlap in APA genes at 4hr and 24hr post-IL-6 treatment and (d) the extent of 3’UTR shortening or lengthening. (e) Correlation between APA genes using either polyAdb or polyAsite annotations. (f) APA and gene expression are not correlated 24hrs post-IL-6 exposure. (g) Only a small number of genes that are significantly alternatively polyadenylated are also significantly differentially expressed.

**Supplemental Figure 3.**
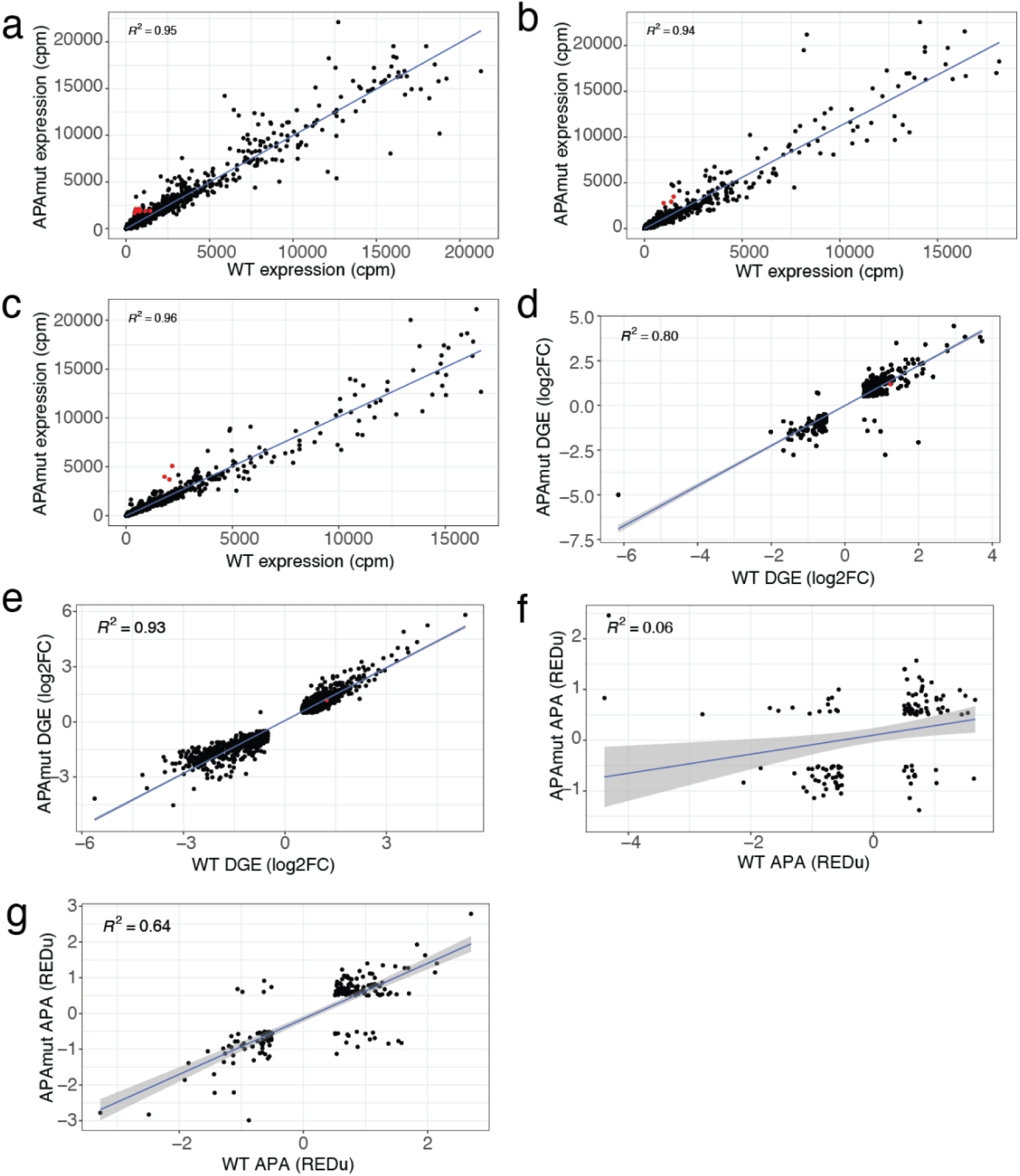
Wild-type and APA-mut HepG2 cells have similar RNA expression and processing. Gene expression is highly correlated in (a) untreated lines, (b) 4 hours post-IL-6 exposure and (c) 24 hours post-IL-6 exposure. Genes identified as significantly differentially expressed (DGE) between untreated and (d) 4 hours IL-6 treatment or (e) 24 hours IL-6 treatment have correlated expression changes in wildtype and APA-mut cells. (f) At 4 hours post-IL-6 treatment alternative polyadenylation (APA) is not correlated between wild-type and APA-mut lines. (g) At 24 hours of IL-6 exposure, genes with differential polyadenylation is correlated between wild-type and APA-mut HepG2 cells.

**Supplemental Figure 4.**
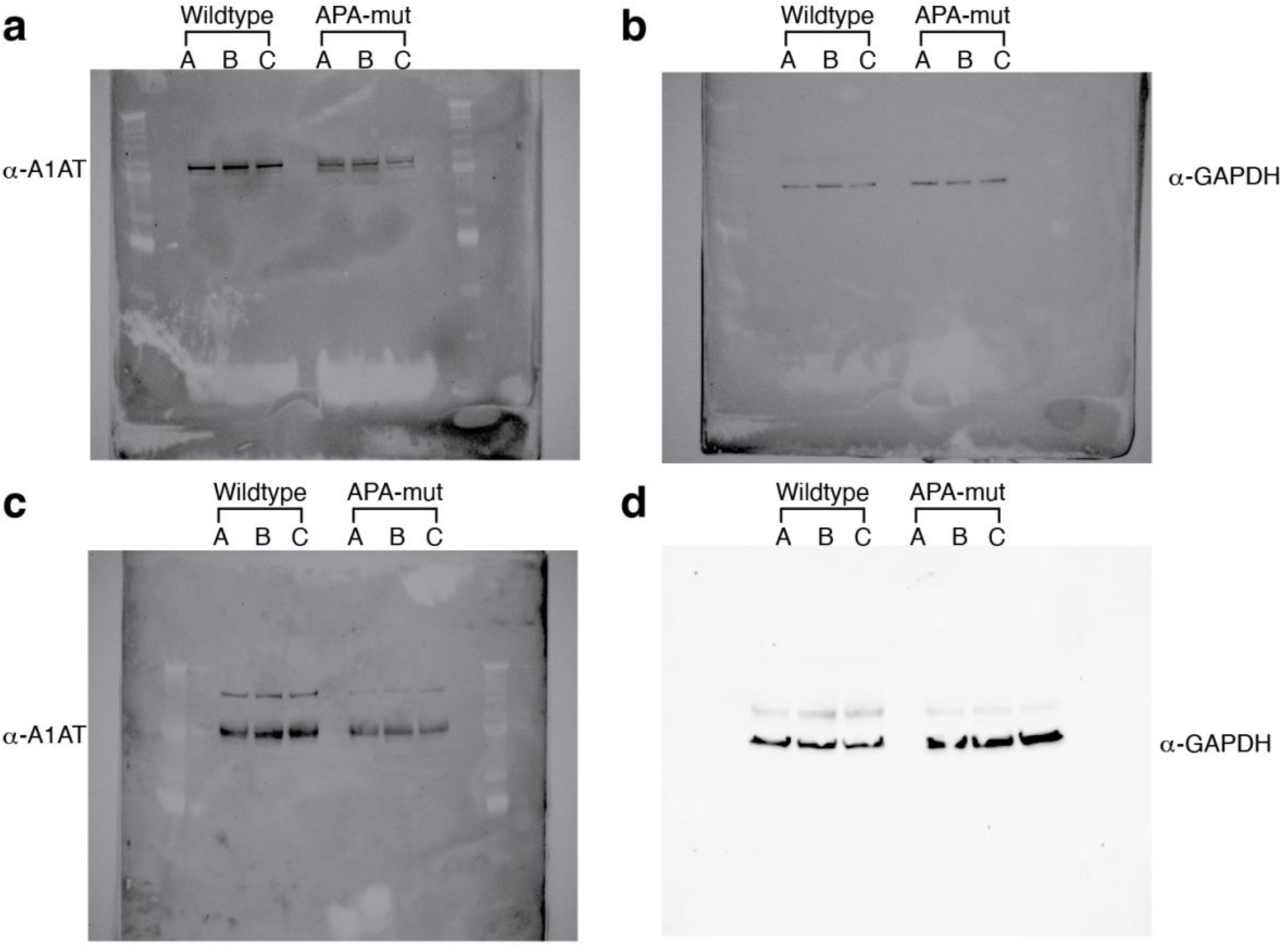
Expression of endogenous A1AT from *SERPINA1* mRNA short and long 3’UTR isoforms. (a,c) A1AT protein levels are decreased in APA-mut CRISPR cells lines compared to wildtype HepG2 cells that predominantly express the long 3’UTR isoform. (b,d) GAPDH was used for normalization.

**Supplemental Figure 5.**
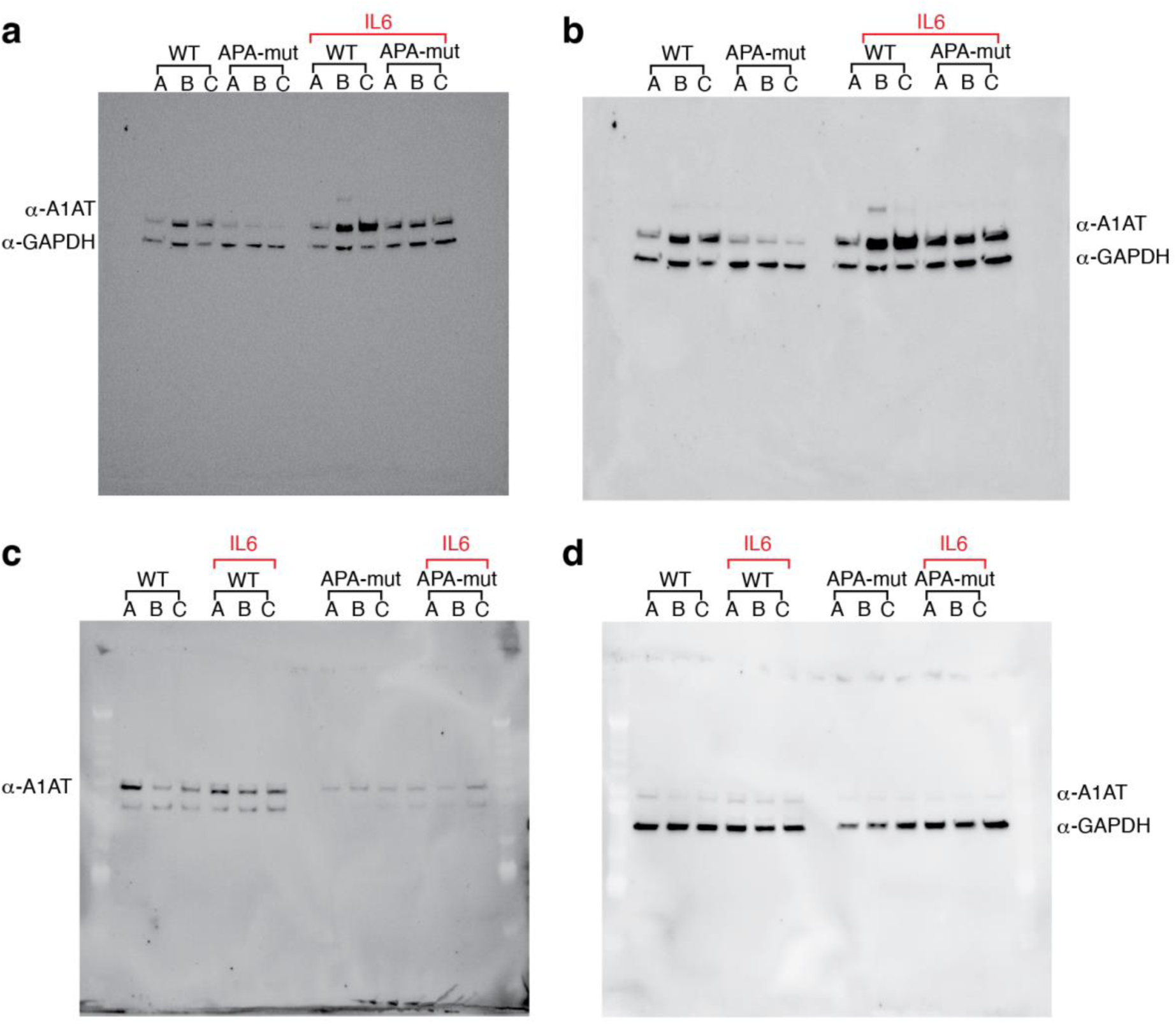
Impact of IL6 treatment on endogenous A1AT protein. (a,c) IL-6 increases the amount of A1AT protein 24 hours after treatment. A1AT protein is decreased in APA- mut CRISPR cells lines in comparison to wildtype HepG2 cells. IL-6 treatment does not increase A1AT protein to the same extent in APA-mut cells. (b,d) GAPDH was used for normalization.

**Supplemental Figure 6.**
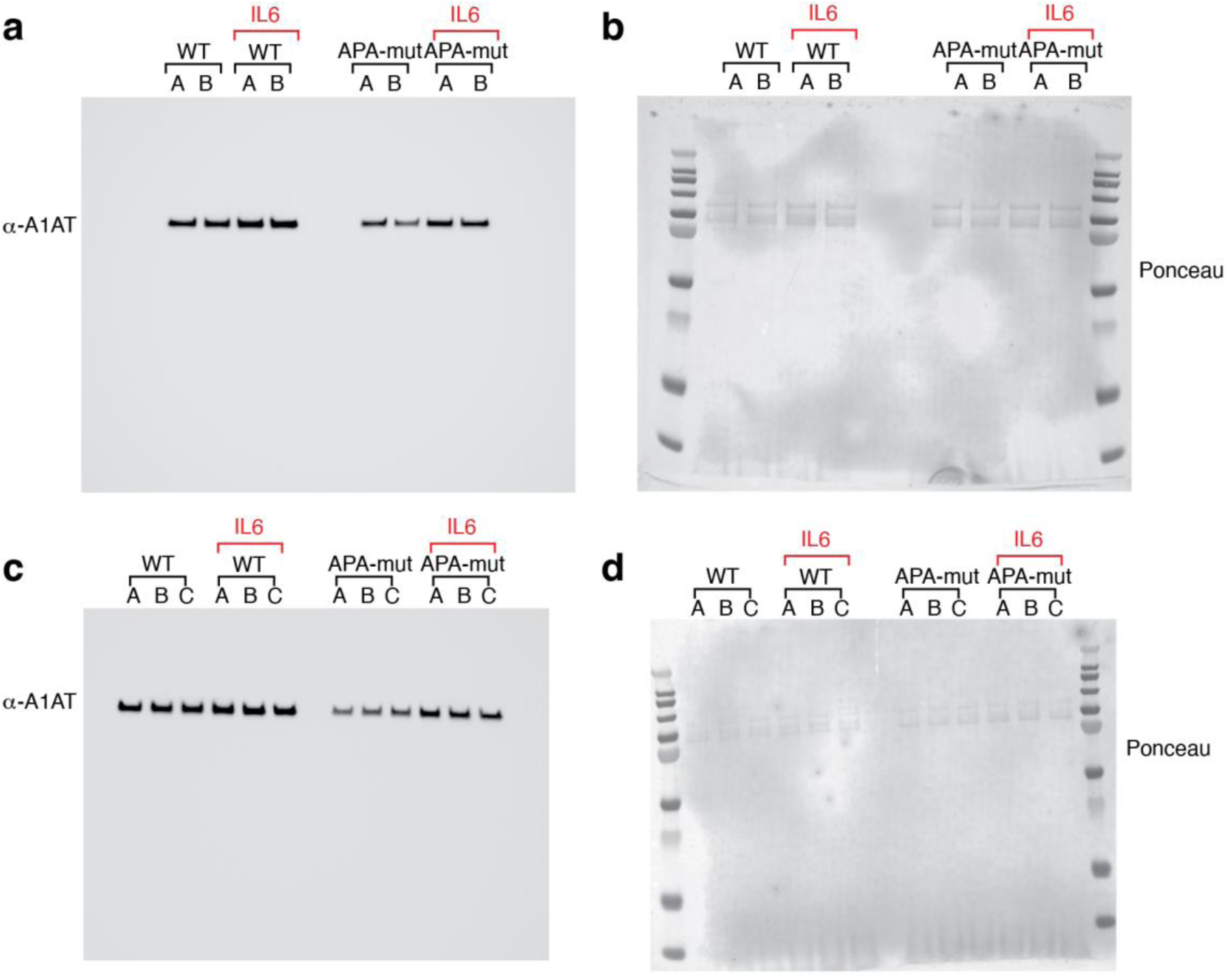
Levels of secreted endogenous A1AT after IL-6 treatment in HepG2 and APA-mut cell lines. (a, c) A1AT protein levels in cell media are decreased in APA-mut CRISPR cells lines with and without IL-6 treatment. (b, d) Media was normalized by volume and Ponceau S staining was used to quantify loading.

**Supplemental Figure 7.**
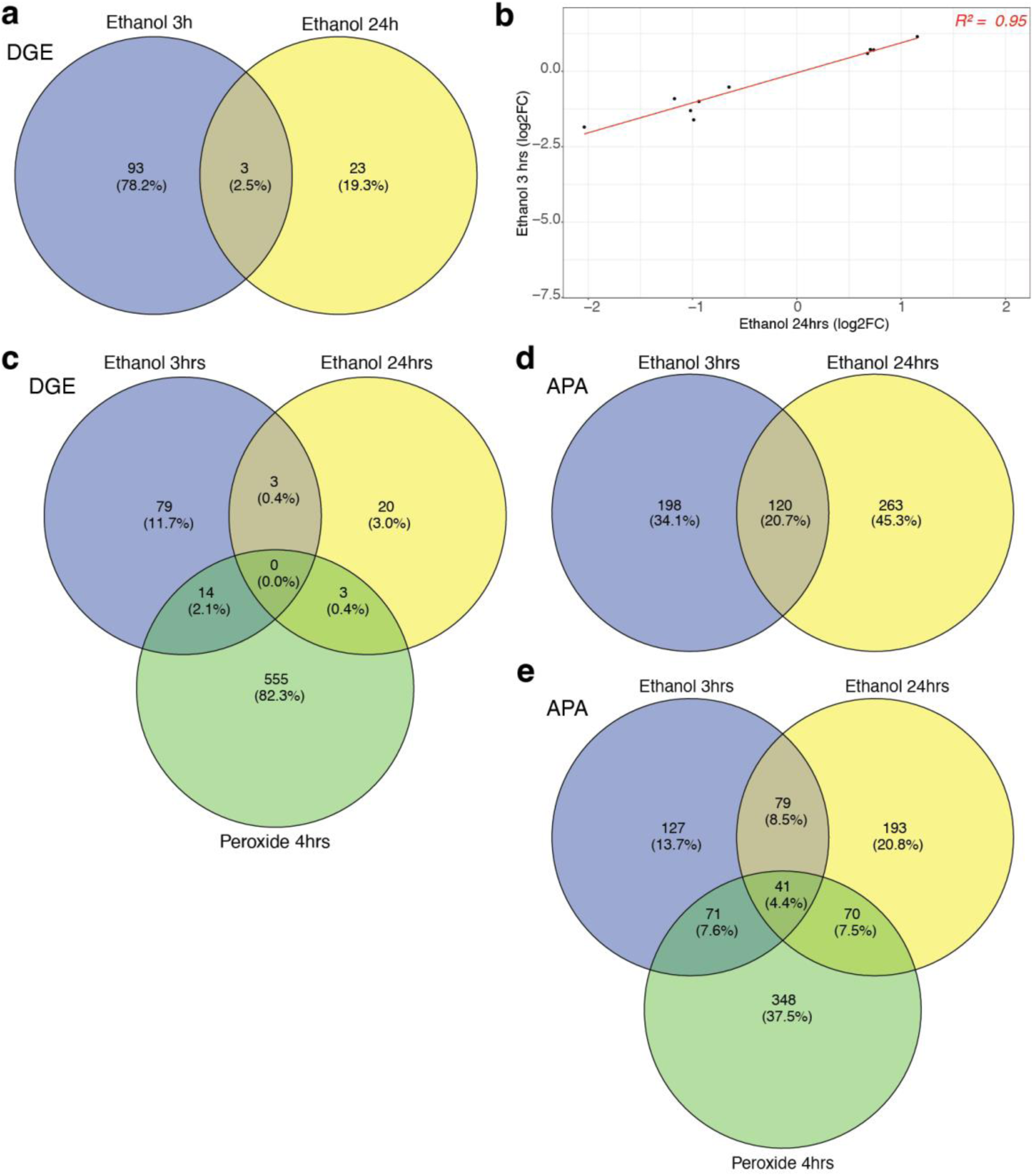
Ethanol and peroxide change gene expression and polyadenylation. (a) Overlap of diffentially expressed genes between 3 and 24 hours of ethanol treatment is minimal but (b) expression is highly correlated for genes affected in both treatments (cutoff log2FC +/- 0.5). (c) More genes are affected by peroxide treatment than ethanol treatment and are not generally shared between treatments. (d) Overlap of significant alternatively polyadenylated genes between 3 and 24 hours of ethanol treatment. (e) Overlap of significant alternatively polyadenylated genes between peroxide treated cells and cells exposed to ethanol for 3 or 24 hours.

**Supplemental Figure 8.**
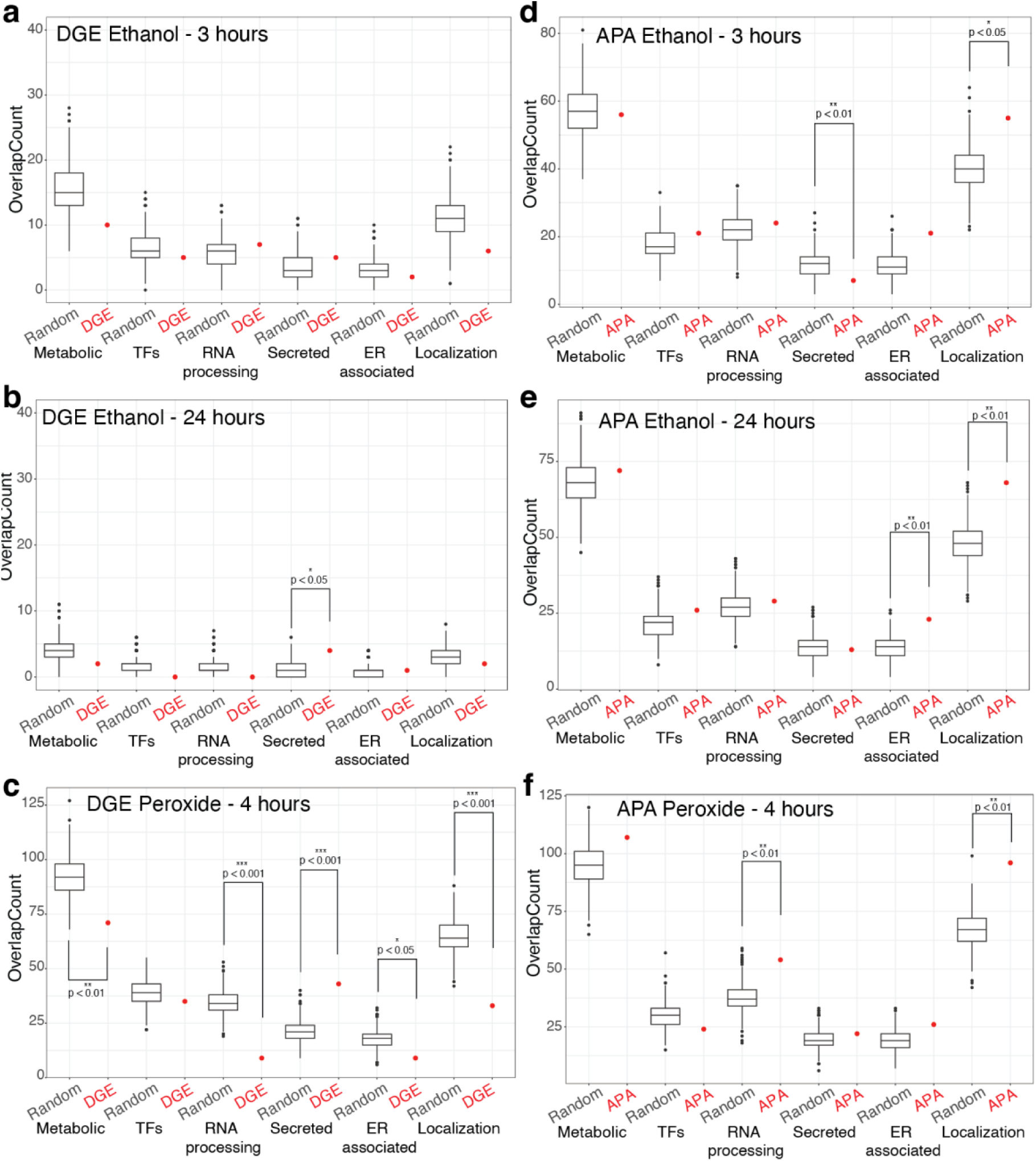
Ethanol has a minor impact of gene expression and 3’ processing compared to peroxide exposure. Select gene lists were tested for overlap between differentially expressed genes after (a-b) ethanol or (c) peroxide exposure compared to randomized expressed genes. The same lists were tested for overlap with alternatively polyadenylated genes after (d-e) ethanol or (f) peroxide exposure. Significance was calculated by permutation test and shown as p<0.001 (***), p<0.01(**) and p< 0.05 (*).

**Supplemental Figure 9.**
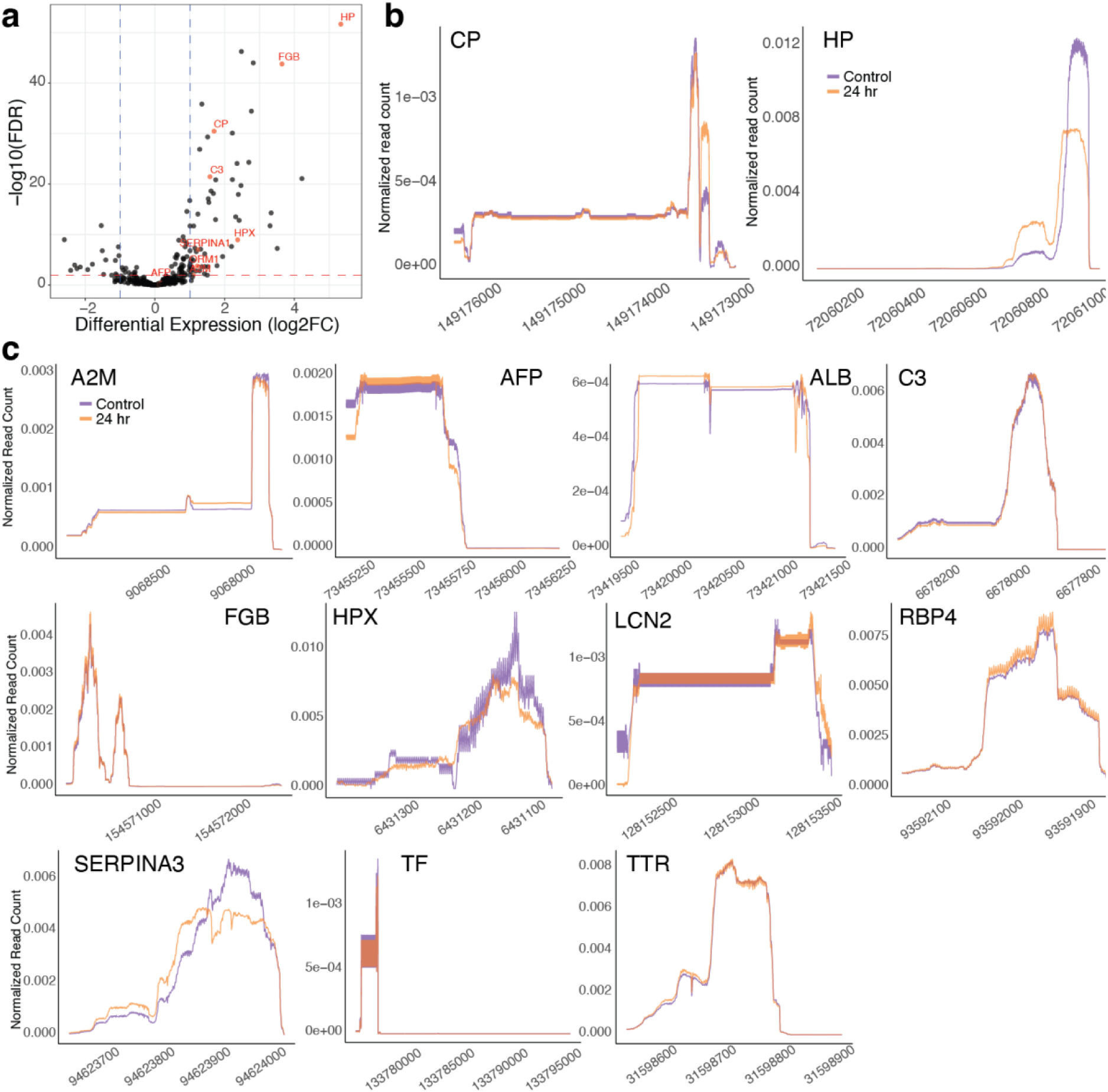
Expression and polyadenylation of acute phase genes. (a) Most acute phase genes are upregulated 24 hours after IL-6 exposure (red) within the group of secreted genes. (b) Ceruloplasmin (CP) and haptoglobin (HP) show altered polyA site selection 24 hours after IL-6 exposure with individual quantification of region normalized read counts across the 3’UTRs. (c) The majority of expressed acute phase proteins do not have differential polyadenylation 24 hours after IL-6 exposure.

**Supplementary Table 1:**
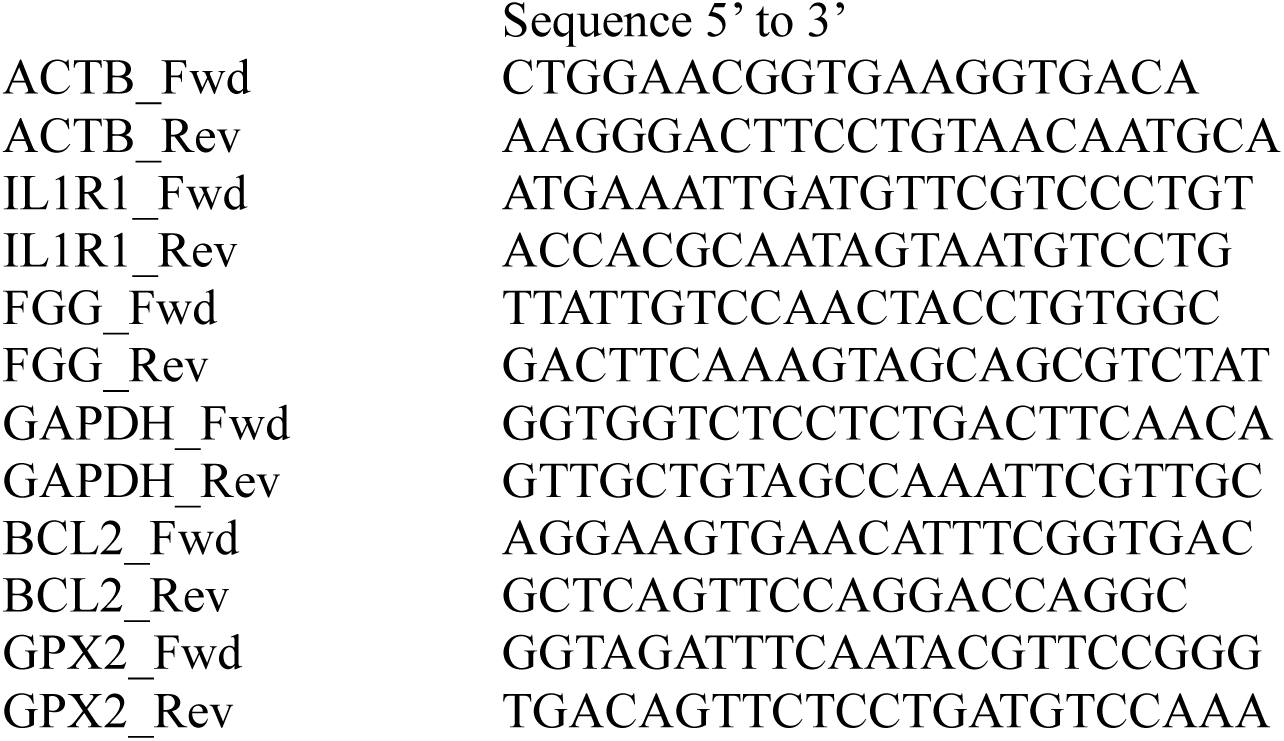
Primers used for qRT-PCR.

